# Oocytes, a single cell and a tissue

**DOI:** 10.1101/2020.02.17.952929

**Authors:** Di Wu, Jurrien Dean

**Author notes:** To whom correspondence should be addressed: Di Wu or Jurrien Dean.

## Abstract

Development of single cell sequencing allows detailing the transcriptome of individual oocytes. Here, we compare different RNA-seq datasets from single and pooled mouse oocytes and show higher reproducibility using single oocyte RNA-seq. We further demonstrate that UMI (unique molecular identifiers) based and other deduplication methods are limited in their ability to improve the precision of these datasets. Finally, for normalization of sample differences in cross-stage comparisons, we propose that external spike-in molecules are comparable to using the endogenous genes stably expressed during oocyte maturation. The ability to normalize data among single cells provides insight into the heterogeneity of mouse oocytes.

## Background

Single cell RNA-seq has revolutionized investigations of transcriptomes that transform cell fate during development. The application of UMI (unique molecular identifiers) now acts as a standard for read quantification, which greatly improves data precision and reproducibility, especially for low count genes [1–3]. Examining the heterogeneity and interactions between cell types benefit the studies of multicellular tissue and tumor development significantly.

As a single cell, the mammalian oocyte is unique in its large size, uniform morphology, high RNA content and wide range of nucleic acid species. During oocyte maturation that immediately follows oocyte growth, the maternal transcriptome is dramatically degraded and reshaped in preparation of fertilization and early development. Since single oocyte RNA-seq was introduced [4], it has become increasingly popular for multi-omic studies in mice. These advances provide greater understanding of developmental and genetic variation as well as detailed information in the co-regulation of transcripts that affect oocyte quality and ageing [5–7]. However, there has been no direct evaluation of single oocyte RNA-seq compared to pooled oocytes, or any discussion of different analytical methods. By pairwise analyses of several published RNA-seq datasets of single and pooled oocytes, we now document the high reproducibility of single oocyte RNA-seq. In addition, by performing UMI based or Picard deduplication, we show that deduplication does not provide significant improvement to single oocyte RNA-seq. Finally, we demonstrate that external spike-in molecules, such as ERCC (External RNA Control Consortium), can effectively account for transcriptome size changes during oocyte maturation. We extracted a group of stably transcribed genes (constGenes) during oocyte maturation and illustrated the similarity of using ERCC and constGenes for cross-stage normalization. This normalization allows greater understanding of the oocyte heterogeneity at the germinal vesicle (GV) stage. ConstGenes or ERCC normalization should become standards for cross-stage comparison in future single oocyte sequencing studies.

## Results and discussion

### Single oocyte RNA-seq has high reproducibility

To estimate the reproducibility of mouse oocyte RNA-seq, we obtained several published datasets from single or pooled oocytes at GV (geminal vesicle) and MII (meiosis II) stages which are the beginning and ending of oocyte maturation, respectively [6–10]. We processed all raw sequencing files in parallel. Due to the different distribution of transcripts from all the samples, we only took the coding reads for comparison (Fig. 1a; Table S1). Interestingly, single oocyte RNA-seq (GSE141190, GSE96638, GSE44183) had higher correlation within the group than when compared to pooled oocytes of the same stage (Fig. 1b). In addition, MII stages exhibit higher deviation of poly(A) vs RiboMinus sequencing results, possibly due to the existence of dormant RNAs and hyper polyadenylated mRNAs being actively translated (Fig. 1b) [11]. By principal component clustering, single oocyte groups show higher similarity, though experiments/methods dominate the difference (Fig. 1c-d). Thus, we concluded that single oocyte sequencing can generate highly reproducible and consistent results.

**Figure 1.**
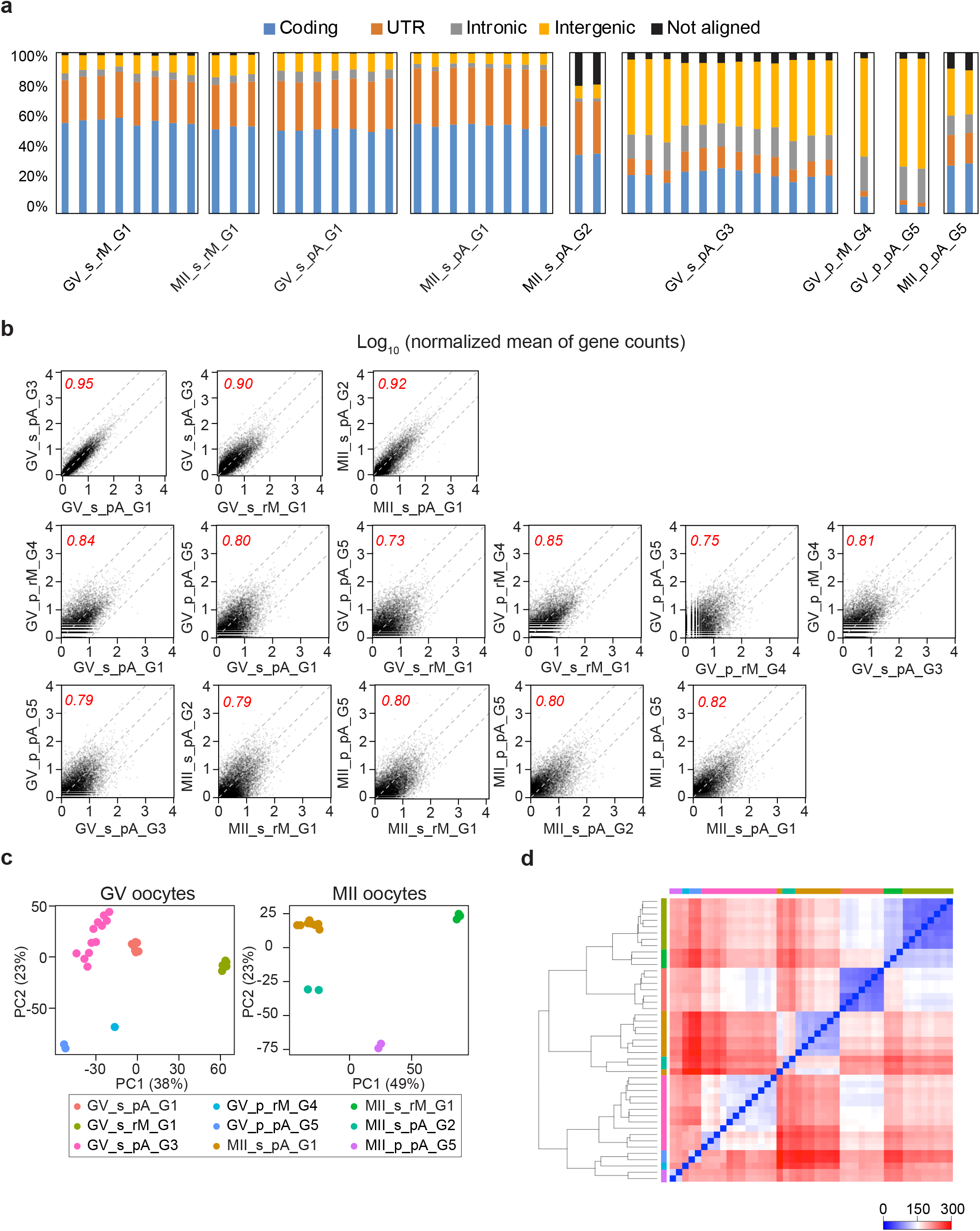
Single mouse oocyte RNA-seq exhibited higher reproducibility. **a** Bar graph of read coverage in coding, UTR, intronic and intergenic regions in different datasets. **b** Plots of transcript abundance regression analysis of single or pooled oocytes RNA-seq results. Normalized mean of gene counts: mean of gene counts from all libraries normalized by library sizes in each condition. In each plot, the x-axis and y-axis are the log_10_ count values from different sequencing results. Three gray dashed lines: y=x, y=x+1 and y=x−1. Red number: the correlation coefficient of each comparison. The library names: GV/MII_single/pooled_polyA/riboMinus_GEO abbreviation. The GEO abbreviation is listed in *Availability of data and materials*. **c-d** Principal component analysis (c) and sample distance matrix (d) of the single and pooled oocytes RNA-seq datasets at GV and MII stages.

### Deduplication improved the single oocyte RNA-seq limitedly

Single cell RNA-seq is susceptible to a range of biases, including gene capture, reverse transcription and cDNA amplification [12]. The incorporation of UMI (unique molecular identifiers) significantly improves single cell sequencing reproducibility by quantifying reads with more precision [3]. To test whether UMI also benefits single oocyte RNA-seq, we re-analyzed the GSE141190 RiboMinus RNA-seq results using UMI quantification (Fig. 2a-b) which determines duplicates by both UMI and insert reads. The N8 UMI, capable of distinguishing up to 65,536 molecules, is sufficient to distinguish the ~20,000 different RNAs expressed in mouse oocytes [3, 7]. On average, the number of reads of the Dedup (UMI-based) samples were 43%±15% of their Original (without deduplication) samples (Fig. 2c-d; Table S2). After filtering out low-count genes, the linear regression of gene counts in Original and Dedup groups, normalized by library size, exhibited high correlation at the same stage (Fig. 2e).

**Figure 2.**
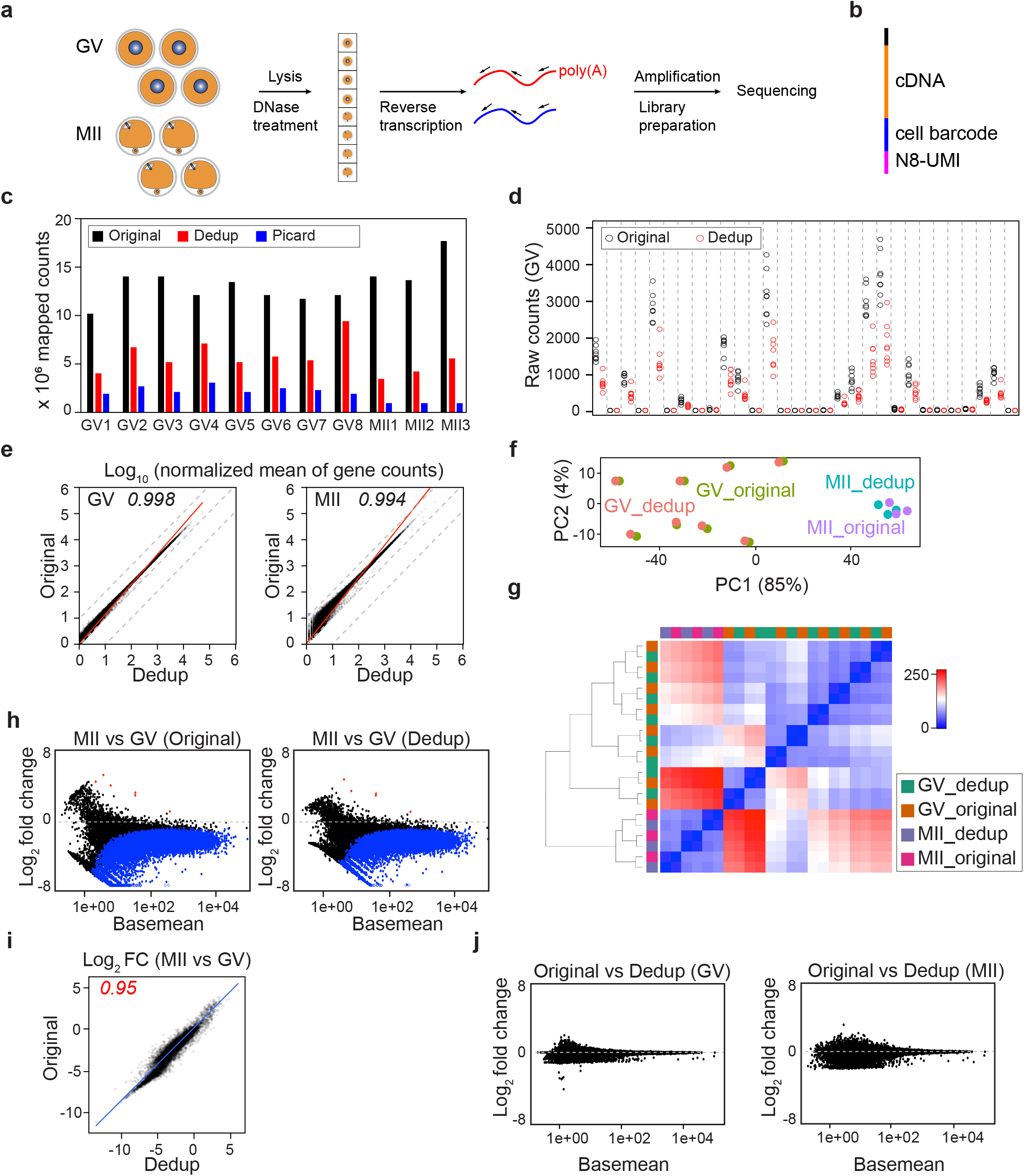
Deduplication of single oocyte RNA-seq reads does not significantly improve results. **a** Outline of single oocyte RNA-seq adapted from Ovation Solo RNA-seq system. **b** Structure of the sequencing reads, including the insert (cDNA), cell barcode and N8 random sequence (UMI, unique molecular identifier). **c** Bar graph of Original, UMI-based (Dedup) and Picard deduplicated (Picard) counts. **d** Dot plot of the Original and UMI-based (Dedup) gene counts at GV stage. The chosen genes are the top 30 after ranking by their Ensembl gene ID. **e** Plots of transcript abundance regression analysis of Original vs Dedup gene counts normalized by library sizes. **f-g** Principal component analysis (f), sample distance matrix (g) of combined Original and Dedup samples. **h-i** Differential analysis of MII vs GV in Original and Dedup groups (h) and the comparison of the log_2_FC (MII vs GV) in Original and Dedup groups (i). **j** MA plots of differentially expressed genes of Original vs Dedup in GV and MII oocytes.

In addition, we performed downstream differential analysis using both Original and Dedup counts. The Original and Dedup counts from the same oocyte were similar, documented both by hierarchical clustering of principal components and distance calculations (Fig. 2f-g). The MII vs GV differential analysis in both Original and Dedup groups also were very similar (Fig. 2h-i; Table S3). On the other hand, no significant change in gene expression was identified (*P*-adjusted < 0.01) when comparing Original and Dedup at the same stage (Fig. 2j; Table S3) which suggests that UMI deduplication does not significantly improve the quantitative precision in single oocyte RNA-seq.

To further evaluate how important deduplication is to single oocyte RNA-seq, we took advantage of Picard deduplication, which defines duplicates based only on mapping coordinates. As expected, Picard trimmed off even more reads (Fig. 1c). Nevertheless, the correlation between Original and Picard remained high though obvious deviation were present due to trimming of reads (Fig. 3a). As expected, differential analyses of Original vs Picard at the GV or MII stages documented that a certain group of genes, including those encoding ribosomal proteins (*Rpl19*, *Rpl32*, *Rps20*) were sensitive to deduplication (Fig. 3b; Table S4). We also used the ERCC (External RNA Controls Consortium) spike-in molecules to visualize the linearization between their counts and raw concentrations. All generated a high correlation of the ERCC molecules compared with their raw concentration (Fig. 3c-e). Thus, we conclude that deduplication provides only a limited advantage to the robustness of single oocyte RNA-seq, possibly due to the high RNA abundancy that makes the oocyte more like a tissue rather than a standard single cell.

**Figure 3.**
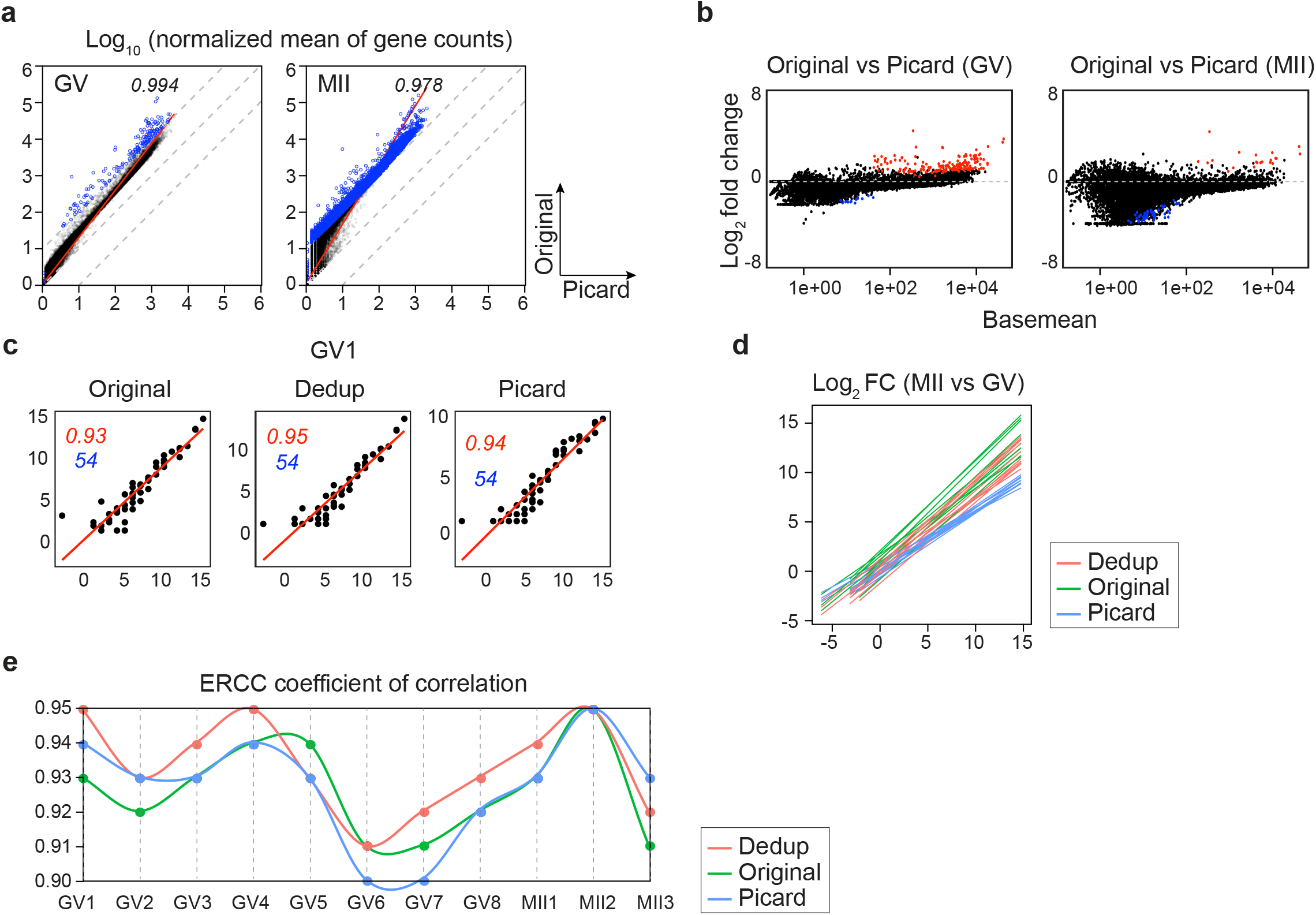
Picard deduplication of single oocyte RNA-seq reads does not significantly improve results. **a** Plots of transcript abundance regression analysis of Original vs Picard gene counts normalized by library sizes. **b** MA plots of differentially expressed genes of Original vs Picard in GV and MII oocytes. **c-e** ERCC regression between the sequenced counts to their original concentration at GV and MII stages from the Original, Dedup and Picard groups. An example of ERCC molecules linearization is in (c), the summary of all groups is in (d) and the coefficients of correlation are summarized in (e).

### External and internal normalization for cross-stage comparison

It has been long known that the transcriptome undergoes dramatic decrease during mammalian oocyte maturation [7, 10, 13]. Presumably, identification of the decreased and stable RNA in this process could add understanding to the molecular regulation of oocyte development and quality control. Several RNA-seq readouts have been used to represent cross-stage differences such as FPKM (Fragments Per Kilobase Million) and RPM (Transcripts Per Kilobase Million) which are normalized by gene length and library size [10, 14]. However, this library-size based quantification indicated as many genes with increased abundance as with decreased abundance which is difficult to explain biologically in the absence of transcription during oocyte maturation. Gfp/Rfp spike-in molecules have also been used to quantify changes in the transcriptome size, but not for differential analysis of individual genes [10]. Here, we propose that the ERCC spike-in mix can serve as control genes for single oocyte RNA-seq. We performed differential analyses using ERCC-dependent and median ratio dependent normalization (GSE141190, GSE96638). Within the same stage (GV), median ratio normalization (ERCC-independent) and the ERCC normalization produced roughly similar results, although ERCC normalization had more within-group deviation (Fig. 4a). However, ERCC normalization dramatically altered the results in cross-stage comparison in the GSE141190 dataset [7]. The absence of genes showing increased abundance is consistent to the absence of transcription during oocyte maturation, suggesting that external spike-in molecules provide better normalization for libraries with variated transcriptome size [7].

**Figure 4.**
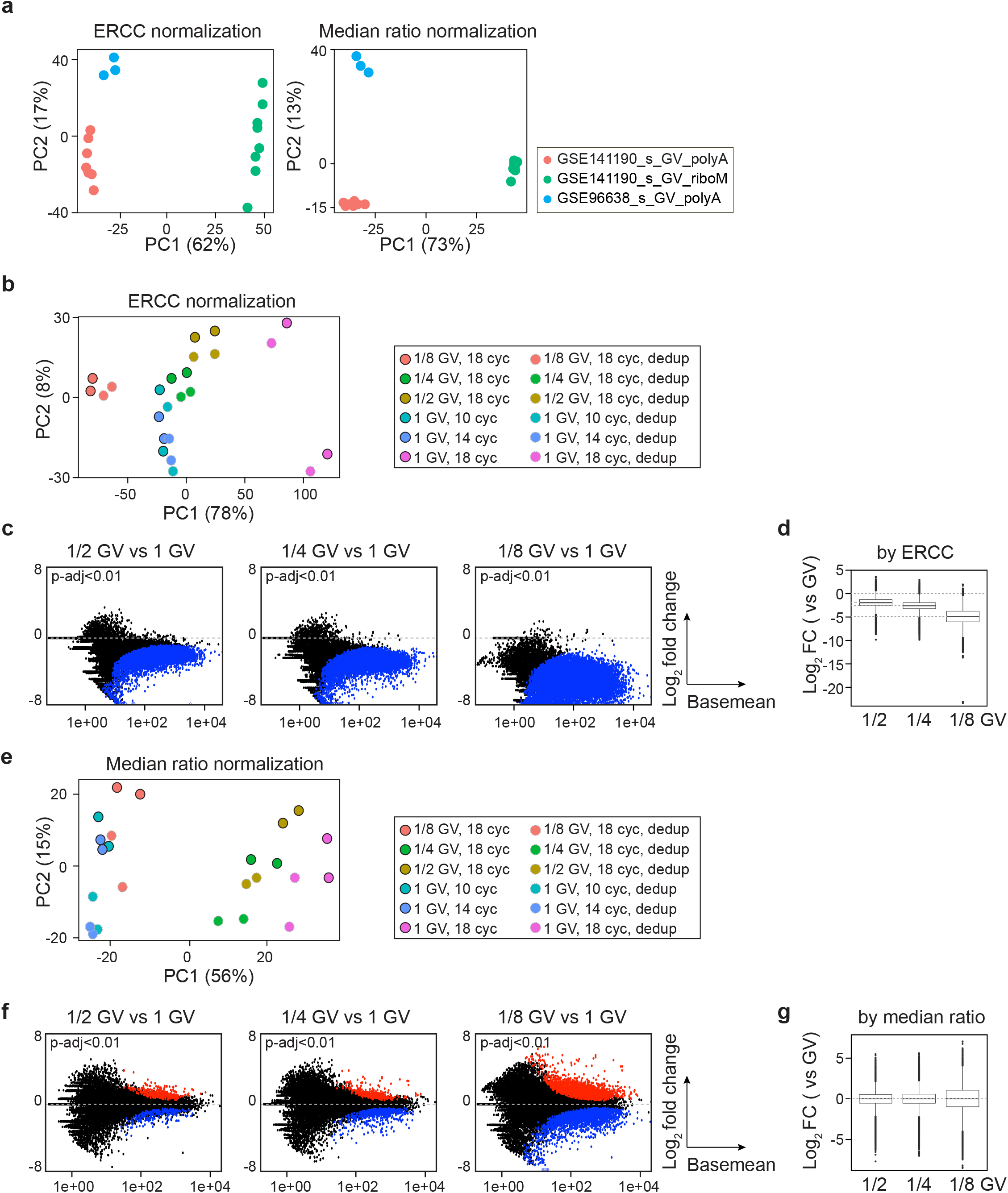
ERCC (External RNA Controls Consortium) spike-in molecules can normalize transcriptome size difference between samples. **a** Principal component analysis (PCA) of ERCC-containing samples, including GSE141190 and GSE96638, by ERCC or Median ratio normalization. **b-g** ERCC or median ratio normalization of GV oocyte fractions (1/2, 1/4, 1/8) and different amplification cycles (10, 14, 18). **b, e** PCA of GV fractions and different amplification cycles by ERCC or median ratio normalization. **c, f** MA plots showing the differentially expressed genes compared between oocyte fractions by ERCC or median ratio normalization. **d, g** Summaries of log_2_ fold change in (c) and (f).

To better illustrate ERCC normalization, we performed poly(A) GV sequencing using different fractions (1/2, 1/4, 1/8) of oocytes and different amplification cycles during library preparation. The PCA generated from ERCC normalization presented greater similarity between 1/4 GV (18 cycles), 1/2 GV (18 cycles), 1 GV (10 cycles) and 1 GV (14 cycles). The 1/8 GV (18 cycles) and 1 GV (18 cycles) were clustered farther away, presumably representing either less robust library representation or over amplification (Fig. 4b). The differential analysis also confirmed the overall smaller library sizes of the GV fractions compared to the intact GV (Fig. 4c-d; Table S5). On the other hand, the PCA generated from median ratio normalization had a different pattern: 1/8 GV (18 cycles), 1 GV (10 cycles) and 1 GV (14 cycles) were clustered together, while 1/2 GV (18 cycles), 1/4 GV (18 cycles) and 1 GV (18 cycles) were clustered together (Fig. 4e). The differential analyses also revealed no indication of the “overall reduction” in GV fractions (Fig. 4f-g; Table S6). Thus, we conclude that exogenous spike-in accounts for changes in library size and facilitates investigation of transcriptome degradation during oocyte maturation.

To allow cross-stage comparison of sequencing samples when ERCC is unavailable, we have extracted a set of constant genes (constGenes) during oocyte maturation. These 147 genes exhibit little change (less than 50%) in transcript abundance from GV to MII whether obtained by poly(A) RNA-seq or RiboMinus RNA-seq (Fig. 5a; Table S7). The constGenes span a large range of gene length (~0.5-27 kb), have GC content ~30%-60%, contain protein coding genes and lncRNAs (Table S8). As expected, the differential analysis normalized by constGenes were very similar to those normalized by ERCC (Fig. 5b-f; Table S9-10).

**Figure 5.**
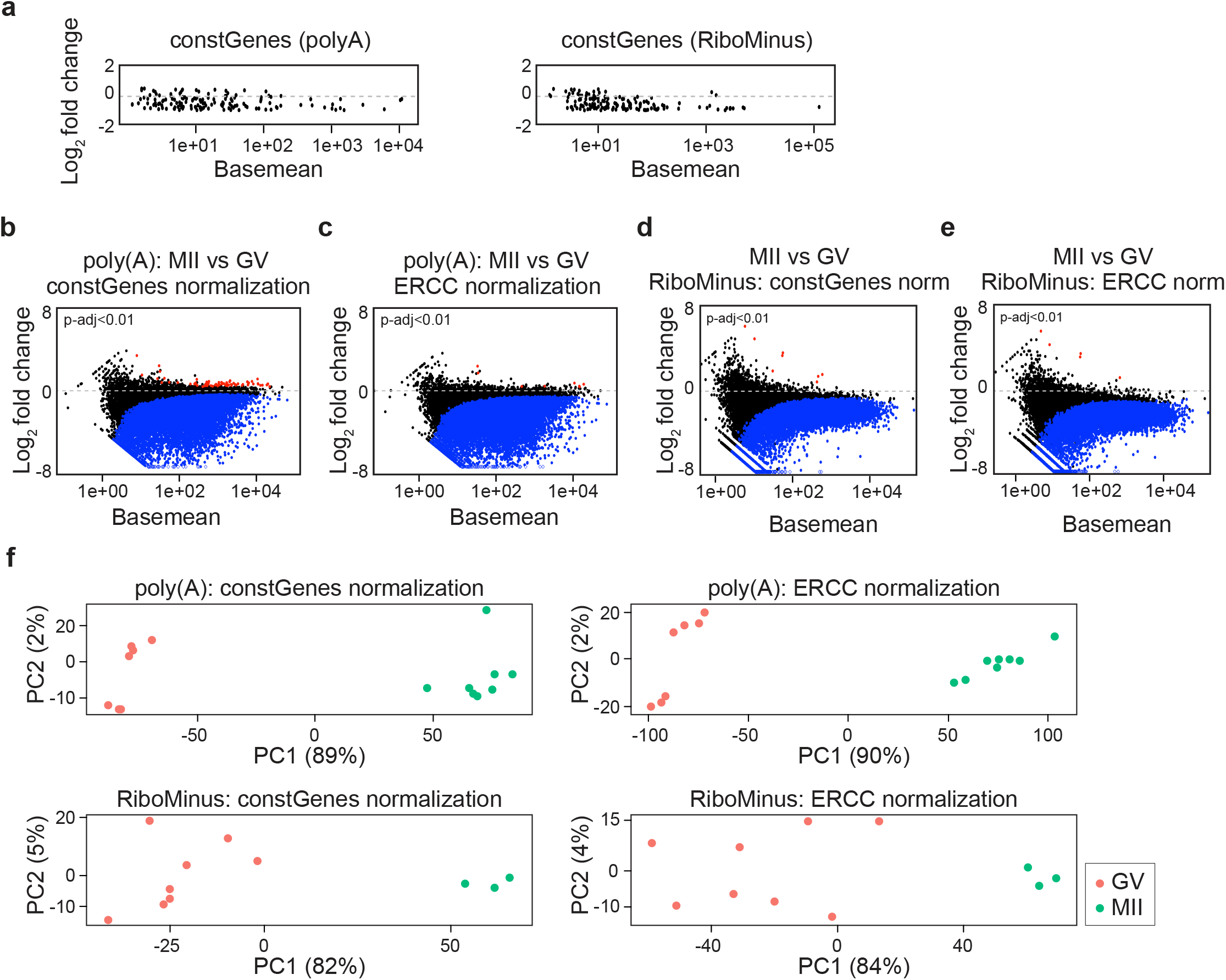
Constant genes (constGenes) can also normalize library differences. **a** MA plots of selected constant genes (constGenes) from poly(A) and RiboMinus RNA-seq results. **b-e** MA plots showing the differentially expressed genes in poly(A) sequencing (b-c) and in RiboMinus sequencing (d-e) from GV to MII stages in GSE141190, normalized by either constGenes (b, d) or ERCC (c, e). Blue/red dots: genes having decreased/increased abundance by *P*-adjusted <0.01. **f** Principal component analysis of GV and MII oocytes from GSE141190 by constGenes or ERCC normalization.

### Better understanding of GV oocyte heterogeneity

Lastly, we investigated the heterogeneity of GV oocytes divided into two subpopulations by nuclear configuration: SN (surrounded nucleolus) and NSN (non-surrounded nucleolus) oocytes [15]. SN oocytes have fully condensed chromatin, no transcription and are more competent for maturation. NSN oocytes have residual transcription and an incomplete meiotic apparatus that reduces developmental competence [15]. The different transcriptome of NSN and SN has been studied by microarray and pooled sequencing which indicated different levels of ribosomal protein and components of the meiotic spindle [16, 17]. Ribosomal RNA sequencing indicated that upregulated 5’ETS of rRNA was linked to the NSN block to maturation [7]. Here, we inspected 3 NSN and 4 SN oocytes by unsupervised clustering and by analysis of the ribosomal RNA reads in the GV samples (Fig. 6a-b). The differential analysis detected a slight down regulation of the transcriptome during the NSN to SN transition (Fig. 6c; Table S11). The differentially expressed genes were enriched with several endomembrane components such as endoplasmic reticulum, Golgi and lysosome that are known to rearrange extensively during nuclear envelope breakdown (Fig. 6d) [18, 19]. Other events such as zinc-finger protein mediated chromatin modification associated with transcriptional inactivation, RNA-binding protein mediated RNA export causing transcript degradation, and abundance of cell division related proteins were also significantly enriched (Fig. 6d) [6, 20].

**Figure 6.**
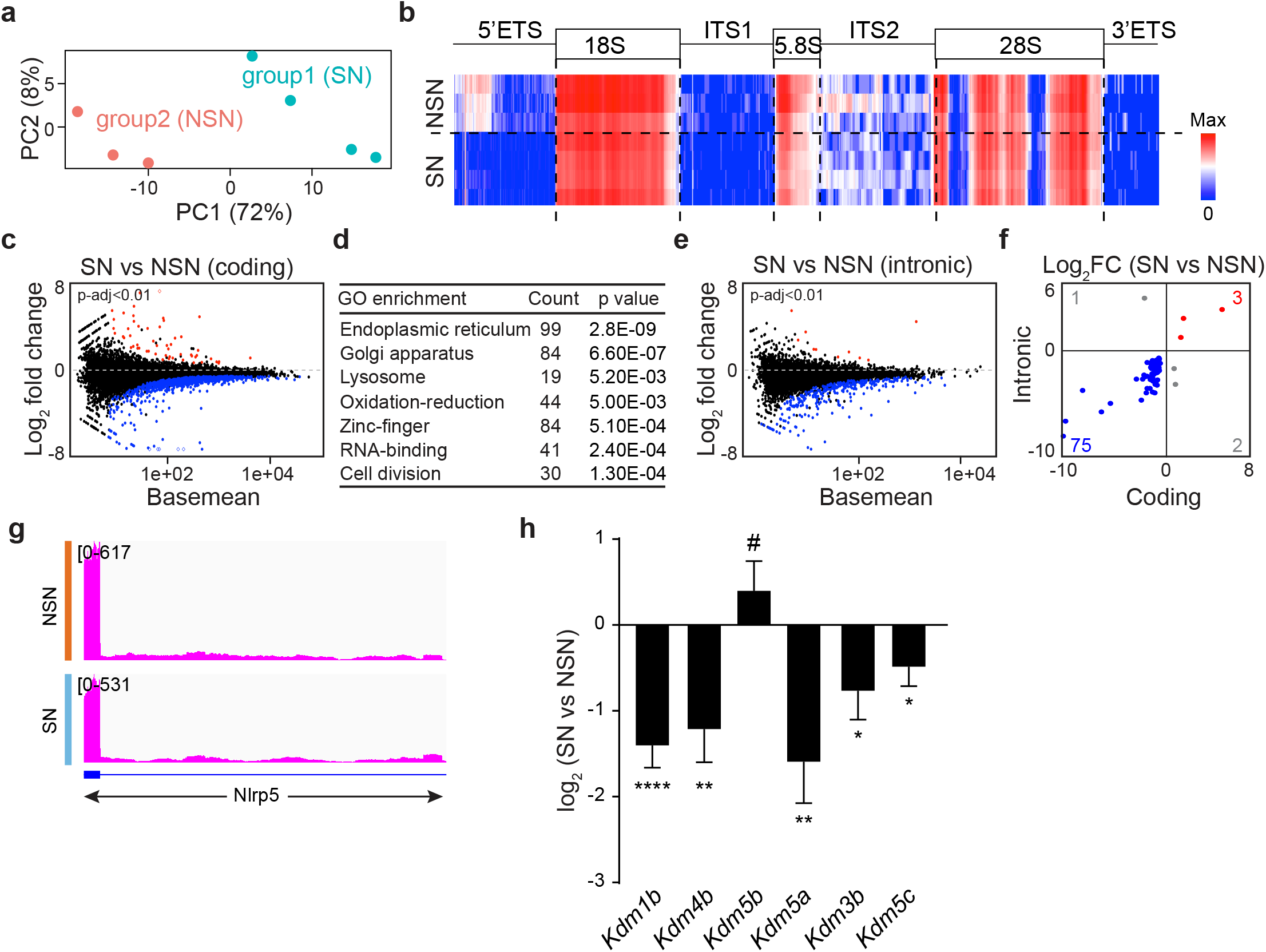
Single oocyte RNA-seq allows better evaluation of oocyte heterogeneity. **a** PCA of GV oocytes from the GSE141190 polyA GV samples. Two groups could be visualized based on the distribution, which were putatively NSN (non-surrounded nucleolus) and SN (surrounded nucleolus) indicated by (b). **b** Heatmap of rRNA coverage obtained from GSE141190 polyA GV samples mapped to the rRNA genomic region. **c-d** MA plots of differentially expressed genes in SN vs NSN GV oocytes counted at coding region (c) and their functional analysis (d). **e** MA plots of differentially expressed genes in SN vs NSN GV oocytes counted at intronic region. **f** Quadrant chart showing coding and intronic sequences that overlapped significantly in differentially expressed genes. **g** Plot from Integrative Genomics Viewer (IGV) showing the read distribution of *Nlrp5* in the SN and NSN oocytes. Blue box: coding region; blue line: intron region. **h** Differential expression fold change of SN vs NSN of the chromatin modification genes.

Interestingly, intronic regions exhibit similar decreases (Fig. 6e; Table S11). The majority (75/81) of the double significantly changed genes shared decrease of coding and intronic reads (Fig. 6f). In addition, the constant distribution of the reads suggested that the residual introns are likely to originate from unspliced pre-mRNAs, which is consistent with the higher transcription activity in NSN oocytes (Fig. 6g). Functional annotation of the 81 differentially expressed genes are significantly enriched in cytoskeleton related genes (modified Fisher Exact *P-value* 0.037). These include *Farp1*/*2*, *Dnm3*, *Eml6*, *Epb41l2*, *Flnb*, *Fmn1*, *Myo10*/*19*, *Arhgef18* and *Sptan1* which provide evidence that SN oocytes contain more proteins required for meiotic spindle organization. Moreover, the known decreased histone demethylase transcripts during the NSN to SN transition were also found to be decreased in the SN group, including *Kdm1b*, *Kdm4b*, *Kdm5a*, *Kdm3b*, *Kdm5c* (Fig. 6h; Table S12) [6].

In summary, based on higher reproducibility and known genetic/developmental heterogeneity, single cell RNA-seq appears to be the best method for transcriptome analyses of mouse oocytes. When an overall change of transcriptome size is anticipated, external spike-in or the constGenes are recommended for normalization to provide better insight into biological changes of the transcriptome.

## Methods

### Oocyte collection and culture

Ovaries were dissected, washed with PBS, and transferred into M2 medium (CytoSpring, M2114) plus milrinone (2.5 μM). The ovaries were pierced mechanically with a 30-gauge needle to release oocytes and only fully-grown oocytes (GV-intact oocytes) were collected for further experiments.

### RNA-seq library preparation of intact and fractions of mouse oocytes

Individual and fractions (1/2, 1/4, 1/8) of mouse oocyte RNA-seq libraries were prepared according to a published pipeline with minor modifications [21]. Briefly, single oocyte at desired stages were collected and transferred individually into 2.5 μl RLT Plus (Qiagen) and stored at −80 °C. The single oocyte lysis was diluted 1:2, 1:4 and 1:8 which represented fractions of oocytes. To prepare libraries for sequencing, 1 μl of the 10^5^-fold diluted ERCC spike-in mix (Thermo Fisher Scientific, 4456740) was added to 2.5 μl of each single or fraction lysis samples.

Poly(A) RNA was isolated by oligo (dT) beads, reverse transcribed, amplified and purified [21]. Individual samples were tested for different amplification cycles (10, 14, 18), and different fractions of oocytes (1/8, 1/4, 1/2) were amplified for 18 cycles. After purification, cDNAs were evaluated by Bioanalyzer 2100 (Agilent). Qualified cDNAs were used to construct sequencing libraries by Nextera DNA Sample Preparation Kits (Illumina). The sequencing was performed by the NIDDK Genomic Core Facility using the HiSeq 2500 Sequencing System (Illumina).

### RNA-seq analysis

Raw sequence reads from each sequencing library were trimmed with Cutadapt 2.5 to remove adapters while performing light quality trimming with parameters “-m 10 -q 20, 20”. Library quality was assessed with FastQC v0.11.8 with default parameters. The trimmed reads were mapped to the Mus musculus GRCm38 genome plus ERCC.fasta using STAR 2.7.2a. Multi-mapping reads were filtered using samtools 1.9. For the Original reads group, the uniquely aligned reads were counted using HTseq 0.9.1 as an unstranded library with default parameters. For the Dedup reads group, before reads counting, the deduplication was performed using the UMI index reads and aligned bam files according to the Ovation Solo RNA-seq manual v4. For the Picard reads group, the bam files were pre-processed by Picard to mark and remove duplicated reads. For each count file, a gene/ERCC was considered valid when it had at least 5 reads in at least 2 libraries.

Differential expression between groups of samples was tested using R v3.5.1 with DESeq2 v1.24.0. When ERCC molecules or constGenes were used for normalization, the counts of the ERCC molecules or constGenes were provided as the controlGenes for estimating the size factors (dds <- estimateSizeFactors(dds, controlGenes= grep(“ERCC”, row.names(filtered)))). Otherwise, the default estimate size factors (median ratio) was used. Functional annotation was performed using the DAVID website. For intronic read analyses, the intron gtf file were produced by BEDTools v2.29.0, and reads were directly counted against the special gtf file. For ribosomal RNA read analysis, the reads were directly counted against rRNA genome file [22].

The regression analysis of the gene expression level and the ERCC molecules was performed by R v3.5.1. The sample distance analysis and gene expression differential analysis were performed with DESeq2 v1.24.0. The mapped reads were visualized in IGV 2.3.97.

## Author information

### Affiliations

Laboratory of Cellular and Developmental Biology, NIDDK, National Institutes of Health, Bethesda, Maryland 20892.

### Contributions

Di Wu, Jurrien Dean conceived the project and wrote the manuscript. Di Wu performed the experiments and bioinformatic analysis.

### Corresponding authors

Correspondence to Di Wu or Jurrien Dean.

## Ethics declarations

### Ethics approval and consent to participate

Mice were maintained in compliance with the guidelines of the Animal Care and Use Committee of the National Institutes of Health, NIDDK-approved animal study protocol.

### Consent for publication

Not applicable.

## Competing interests

The authors declare no competing interests.

## Availability of data and materials

The sequencing data reported in this study has been deposited in the Gene Expression Omnibus website with accession code GSE145283.

The other sequencing data used in this study include:

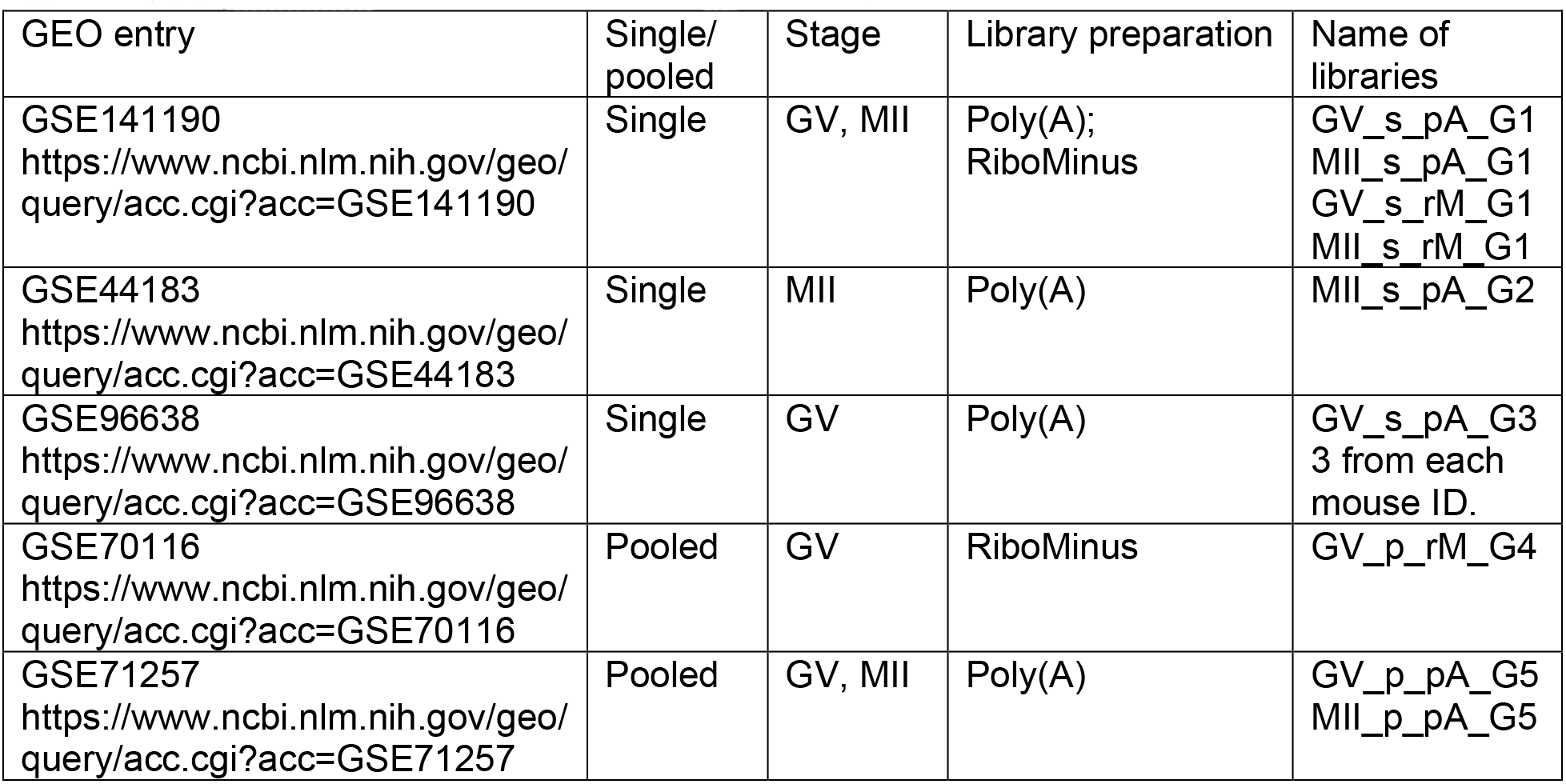

Note that the name of libraries are following the pattern of: ***GV/MII_s***ingle/***p***ooled**_*p***oly***A***/***r***ibo***M***inus**_*G***eo(***1***/***2***/***3***/***4***/***5***)

## Acknowledgements

We appreciate the useful discussions with all the members of the Jurrien Dean lab and with Dr. Ryan Dale, Dr. Guoyun Yu.

## Funding

This work was supported by the Intramural Research Program of NIH, National Institute of Diabetes and Digestive and Kidney Diseases (1ZIADK015603).

## Supplementary information

### Supplementary tables

Table S1. Calculation of the read coverage in all datasets in Fig. 1a.

Table S2. Calculation of Original, Dedup and Picard count results in Fig. 2c-d.

Table S3. Differential analysis results of GV and MII oocytes in Original and Dedup groups in Fig. 2h.

Table S4. Differential analysis results of Original and Picard groups at GV and MII stages in Fig. 3b.

Table S5. Differential analysis results of the GV fractions (1/2, 1/4, 1/8) vs 1 GV through ERCC normalization in Fig. 4c-d.

Table S6. Differential analysis results of the GV fractions vs 1 GV through median ratio normalization in Fig. 4f-g.

Table S7. List of the constGenes obtained from polyA RNA-seq and RiboMinus RNA-seq that remain stable during GV to MII in Fig. 5a.

Table S8. The information of constGenes, including gene length, GC content and description.

Table S9. Differential analysis results of MII vs GV from polyA RNA-seq, normalized by constGenes or ERCC in Fig. 5b-f.

Table S10. Differential analysis results of MII vs GV from RiboMinus RNA-seq, normalized by constGenes or ERCC in Fig. 5b-f.

Table S11. Differential analysis results of the putative SN vs NSN oocytes RNA-seq in Fig. 6c,e.

Table S12. Significantly differential genes in both coding and intronic regions in SN vs NSN RNA-seq in Fig. 6h.

